# Differential Adaptation and Remodeling of the Tumor Microenvironment in Breast Cancer Cells During the Stages of Brain Metastasis

**DOI:** 10.1101/2023.07.13.548875

**Authors:** Xinquan Wu, Yan Zhang, Zhenyong Huang, Ran Li

## Abstract

**Objective:** To integrate macro-transcriptomics and single-cell sequencing to analyze the differential gene expression between primary and brain metastatic breast cancer cells, as well as between active and dormant cancer cells within brain metastases, exploring their adaptation and remodeling abilities at different stages of brain metastasis.

**Methods:** Four public datasets were used: three mRNA microarray datasets from breast cancer and brain metastasis tissues, and one single-cell RNA sequencing (scRNA-seq) dataset from active and dormant brain metastatic cells. Gene differential expression, pathway enrichment, and cell clustering analyses were performed to compare primary and metastatic breast cancer cells, as well as active and dormant cells in brain metastases, focusing on gene expression, metabolic pathways, and functional pathways.

**Results:** Metastatic breast cancer cells showed weakened pathways related to the extracellular matrix and protein digestion. Active cells exhibited enhanced cell cycle regulation, tumor proliferation, and hypoxia resistance pathways compared to dormant cells. Clustering analysis revealed that cluster 6, unique to dormant cells, had enhanced functions in epithelial-mesenchymal transition (EMT), extracellular matrix (ECM), collagen formation, tumor inflammation siganure (TIS), and IL-10 signaling. Cluster 3 and 4 in active cells had enhanced DNA replication and tumor proliferation pathways, respectively.

**Conclusion:** This study highlights the role of cancer cell characteristics and heterogeneity in the invasiveness of brain metastasis. Understanding these mechanisms can guide the development of more effective treatment strategies for breast cancer brain metastasis (BCBM) patients.

## Background

Breast cancer (BC) is one of the most common types of cancer among women worldwide and is also a leading cause of cancer-related deaths in women (1). One of the major challenges in treating BC is managing distant metastasis, which often leads to disease progression and even death. Brain metastasis is a particularly devastating form of metastasis with poor treatment outcomes, resulting in a higher mortality rate (2). A thorough and comprehensive understanding of the underlying biological behavior of brain metastasis is crucial due to the limited treatment options available for patients with brain metastases.

When metastasis begins, BC cells must depart from the primary site, though the molecular mechanisms involved in this process are not fully understood. Some studies suggest a potential association with the tumor microenvironment (TME) (3). After BC cells reach distant organs through the bloodstream or lymphatic system, some cells remain in an active state and form metastases, while others enter a dormant state. The mechanisms underlying this phenomenon are complex and not yet fully elucidated. One hypothesis is that the tumor microenvironment plays a critical role in determining whether cancer cells remain dormant or proliferate. For instance, the recruitment of immune cells to the microenvironment may contribute to tumor cell proliferation and survival (4-6). Recent studies have emphasized the importance of the microenvironment in BC progression, particularly in the context of brain metastasis. For example, one study found significant differences in the tumor microenvironment between brain metastases and primary breast tumors, suggesting that targeted therapies may need to be adjusted to specific microenvironments in the metastatic sites (7, 8). Another study identified key molecular pathways involved in brain metastasis development, including immune signaling and ECM remodeling (8). These findings may serve as potential targets for novel therapies.Moreover, multiple studies have confirmed the presence of cellular heterogeneity within the same lesion of BC, which can result in different functions and behaviors of tumor cells. Some cancer cells within the same tumor may exhibit more aggressiveness and higher invasive and metastatic capabilities (9, 10). After BC cells metastasize to the brain, they need to adapt to the different brain microenvironment compared to the primary site. However, there is limited research on whether cancer cells from the same primary site undergo differentiation in the brain and what characteristics are associated with these differentiated populations. Studies have found that triple-negative BC can be classified into different subtypes based on gene expression patterns, each with distinct clinical outcomes and responses to treatment (11). Therefore, gaining a deeper understanding of the heterogeneity of breast cancer brain metastasis (BCBM) holds positive implications for developing more precise and personalized treatment approaches.

In this study, we utilized four datasets obtained from the Gene Expression Omnibus (GEO, http://www.ncbi.nlm.nih.gov/geo). Among these, GSE209998, GSE125989, and GSE43837 are mRNA-seq datasets containing both breast cancer (BC) tissues and breast cancer brain metastasis (BCBM) tissues. The analysis of these datasets focused on identifying the differences between metastatic and primary lesions. The GSE152818 dataset comprises scRNA-seq data from human BC cell lines metastasized to the mouse brain, serving as an animal model for BCBM. Specifically, dataset GSM4626851 represents proliferating cancer cells from brain metastases, while GSM4626852 corresponds to dormant cancer cells from brain metastases. This portion of the analysis examines the differences between proliferating and dormant cancer cells in BCBM.

By conducting bioinformatics analyses on these two datasets, we compared the differences in metabolic and functional pathways between primary BC cells and metastatic BC cells, as well as between proliferating and dormant cancer cells in BCBM. Additionally, we examined the differences in cell populations between proliferating and dormant cancer cells in brain metastases. By integrating macro-scale transcriptomics with scRNA-seq data, this study explored the potential biological behaviors of BC cells during brain metastasis at two different levels, with the aim of providing valuable insights for the diagnosis and treatment of BC patients.

## Materials and Methods

### Bulk tissue and single-cell sequencing datasets

GEO database is a freely accessible public database. In this study, we utilized four datasets obtained from this database. Among them, GSE209998, GSE125989, and GSE43837 are mRNA-seq datasets containing bulk BC tissues and BCBM tissues, providing a total of 44 brain metastasis samples and 79 primary BC samples. Specifically, GSE209998 consists of 9 brain metastasis samples and 44 primary BC samples; GSE125989 consists of 16 brain metastasis samples and 16 primary BC samples; GSE43837 consists of 19 brain metastasis samples and 19 primary BC samples. The raw data have undergone batch correction during the upload process.

The GSE152818 dataset is a scRNA-seq dataset from a mouse model of human BCBM. In this model, the BC cell line HMT-3522-T4-2 was injected into the left ventricle of four 6-8-week-old female mice under ultrasound guidance. The mice were euthanized one month after injection, and brain dissection was performed to obtain samples. For detailed processing methods, please refer to the sample descriptions (https://www.ncbi.nlm.nih.gov/geo/query/acc.cgi?acc=GSM4626851; https://www.ncbi.nlm.nih.gov/geo/query/acc.cgi?acc=GSM4626852). Dataset GSM4626851 represents proliferating BC cells from brain metastases, while dataset GSM4626852 represents dormant BC cells from brain metastases.

### Data Preprocessing

First, the GSE209998, GSE125989, and GSE43837 datasets were subjected to data cleaning. After obtaining the expression matrix, we removed rows (genes) with an NA value proportion greater than 50% and columns (samples) with an NA value proportion greater than 50%. Then, the impute.knn function from the impute package (version 3.17) in R software (http://www.r-project.org/, version 4.3) was used for missing value imputation, with the Number of neighbors set to 10 for data completion.

After downloading the raw data of GSM4626851 and GSM4626852, the Seurat package (version 4.3) in R software (https://satijalab.org/seurat/) was used to convert the scRNA-seq data into seurat objects and filter out low-quality cells. Following the recommended criteria in the Seurat tutorial based on the number of unique genes per cell (**Figure 1**), cells meeting any of the following conditions were defined as low-quality cells in this study: (1) expressing fewer than 200 genes, (2) expressing more than 8000 genes, or (3) having 5% or more unique molecular identifiers (UMIs) mapped to mitochondrial genes. Finally, the ScaleData function from the Seurat package was applied for batch normalization.

**Figure 1.**
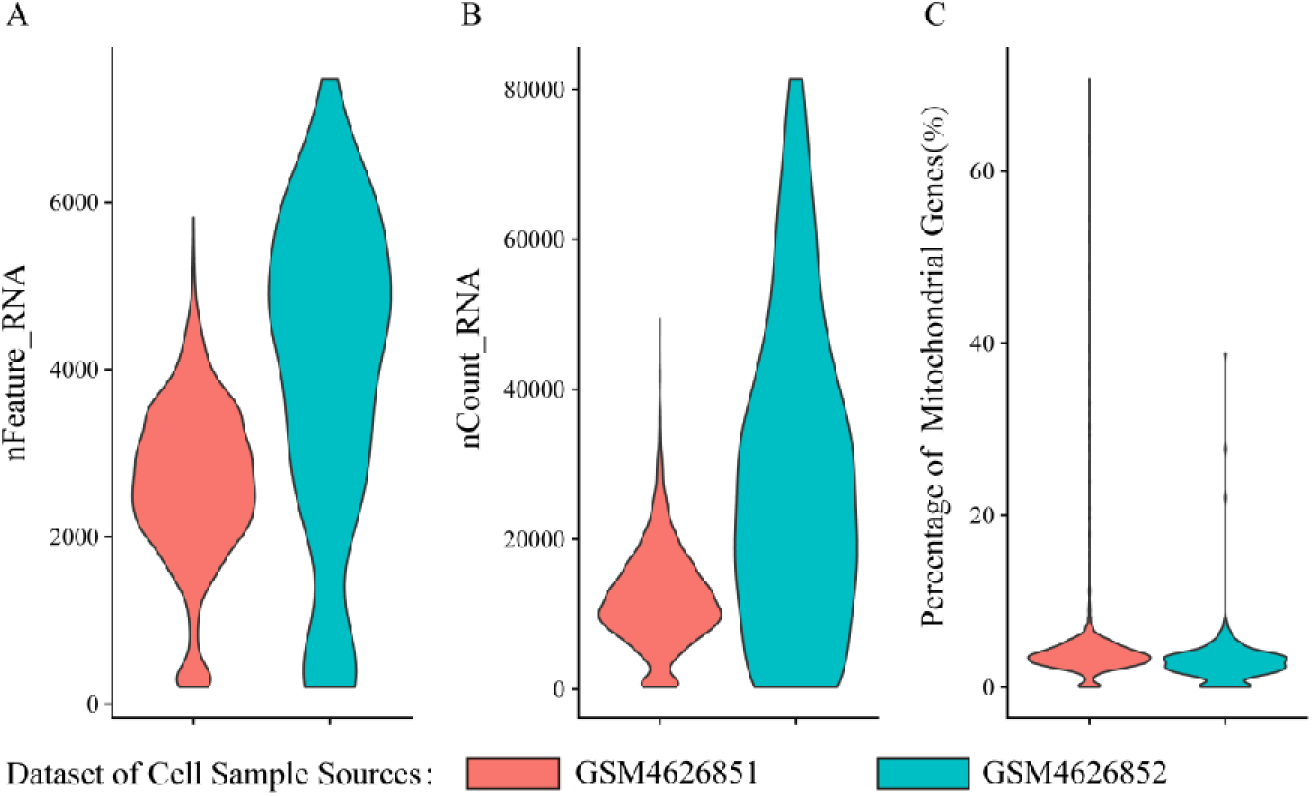
Plots depicting the number of genes detected in each cell (nFeature_RNA) (A), the number of unique RNA molecules detected in each cells (nCount-RNA) (B), and the percentage of reads mapping to the mitochondrial genome (percent.mt) (C).

### Screening of Differentially Expressed Genes

We used the limma package in R software (http://bioconductor.org/packages/release/bioc/html/limma.html, version 3.56) to identify differentially expressed genes (DEGs) between tumor tissues and normal tissues. Fold change (FC) represents the magnitude of gene expression differences. A log |FC| > 1.5 and a p-value < 0.05 were set as the cutoff criteria for determining DEGs. After performing the calculations, differential gene sets were obtained for three datasets: GSE209998, GSE125989, and GSE43837. To visualize the differential results, we used the ggplot2 package in R software (https://ggplot2.tidyverse.org/, version 3.4.1) to create a volcano plot and a heat map, where specific genes are marked.

### Perform Robust Rank Aggregation (RRA) analysis on the datasets of DEGs

The three datasets we used were generated under different experimental conditions and utilized different microarray platforms, making it unreliable to merge and analyze the three gene sets together (12). The principle of Robust Rank Aggregation (RRA) (13) was to compare the actual rankings of each dataset with random order null hypotheses and filter DEGs by assigning significance scores to each gene. In this study, RRA calculated the significance scores for all genes by combining the p-values and logFC values from the differential gene lists of the three datasets. The RRA package (https://cran.r-project.org/web/packages/RobustRankAggreg/index.html, version 1.2) was used for meta-analysis of the ranked gene lists. After adjusting for false discovery rate (FDR), a log |FC| > 1 and a p-value < 0.05 were chosen as the threshold for selecting commonly DEGs. We selected the top 10 upregulated and top 10 downregulated genes (a total of 20 genes) with the highest significance to generate a heatmap and proceeded with further analysis.

### Perform pathway analysis on the commonly upregulated/downregulated genes

The purpose of gene functional enrichment analysis was to assist researchers in understanding the biological roles of a set of genes. In this study, pathway enrichment analysis was performed using background gene sets from the Hallmark, Kyoto Encyclopedia of Genes and Genomes (KEGG), and Gene Ontology (GO) gene sets in the MSIGDB database (14). The R package clusterProfiler (https://bioconductor.org/packages/release/bioc/html/clusterProfiler.html, version 4.8) was used to conduct GO, KEGG, and Hallmark pathway analyses on the top 10 significantly upregulated and downregulated genes obtained from RRA. The cutoff criteria for pathway analysis were adjusted false discovery rate (FDR) with a p-value < 0.05 and a pathway gene count ≥ 3.

### Perform principal component analysis (PCA) and dimensionality reduction on the preprocessed single-cell data

For batch-corrected scRNA-seq data, the FindVariableFeatures function from the Seurat package was used to select highly variable genes. The top 2000 genes with the highest standard deviation were defined as highly variable genes. The RunPCA function from the Seurat package was then utilized to perform PCA on the highly variable genes, calculating the top 20 principal components. Subsequently, the JackStraw plot function from the Seurat package was used to visualize the distribution of p-values for each principal component. The p-value distribution was compared to a uniform distribution (dashed line), and significant principal components were defined as those above the dashed line with statistically significant p-values (P < 0.05). Important principal components are retained for visualization and clustering. Next, the FindNeighbors and FindClusters functions from the Seurat package were employed for cell clustering (resolution 0.5). Finally, the RunUMAP function from the Seurat package was used to perform UMAP (15), a nonlinear dimensionality reduction technique, for visualization purposes.

### Marker genes identification and cell types clustering

The FindAllMarkers function from the Seurat package was used to identify marker genes in each principal component. A cutoff of |log2FC| > 1 and adjusted p-value < 0.05 was applied, and the top 10 marker genes with the highest rankings in each subcluster were selected for cell type clustering.

### Cell type and metabolic pathway analysis of proliferating tumor cell population and dormant tumor cell population

Based on the grouping and cell clustering data, a stacked bar plot of the proportions of different cell groups in each group was generated using the ggplot2 package in R software (https://cran.r-project.org/web/packages/ggplot2/index.html, version 3.4.2). The AverageExpression function was used to calculate the average expression values of genes in the cells included in different groups. Then, the Gene Set Variation Analysis (GSVA) algorithm was applied to load the MSigDB database for differential analysis at the gene set level between different groups. We performed one calculation based on the grouping of cells according to their sources (proliferating cells and dormant cells), and then we performed another calculation by re-grouping the cells based on their clusters. A heatmap was created using the pheatmap package (CRAN.R-project.org/package=pheatmap, version 1.0.12) based on the score of each group for the corresponding pathways.

## Results

### Identification of DEGs between BCBM and primary tumor and RRA analysis of DEGs

Based on the cutoff criteria, we identified 7,974 DEGs in GSE209998, including 2,038 upregulated genes and 5,936 downregulated genes. In GSE125989, we identified 979 DEGs, with 307 upregulated genes and 672 downregulated genes. Additionally, in GSE43837, we identified 1,832 DEGs, including 519 upregulated genes and 1,313 downregulated genes. The volcano plots (**Figure 2A-C**) display these DEGs, while the heatmaps (**Figure 2D-F**) show the top 10 upregulated and top 10 downregulated genes. After RRA analysis on the three DEGs sets, 32 commonly upregulated genes and 158 commonly downregulated genes were identified.

**Figure 2.**
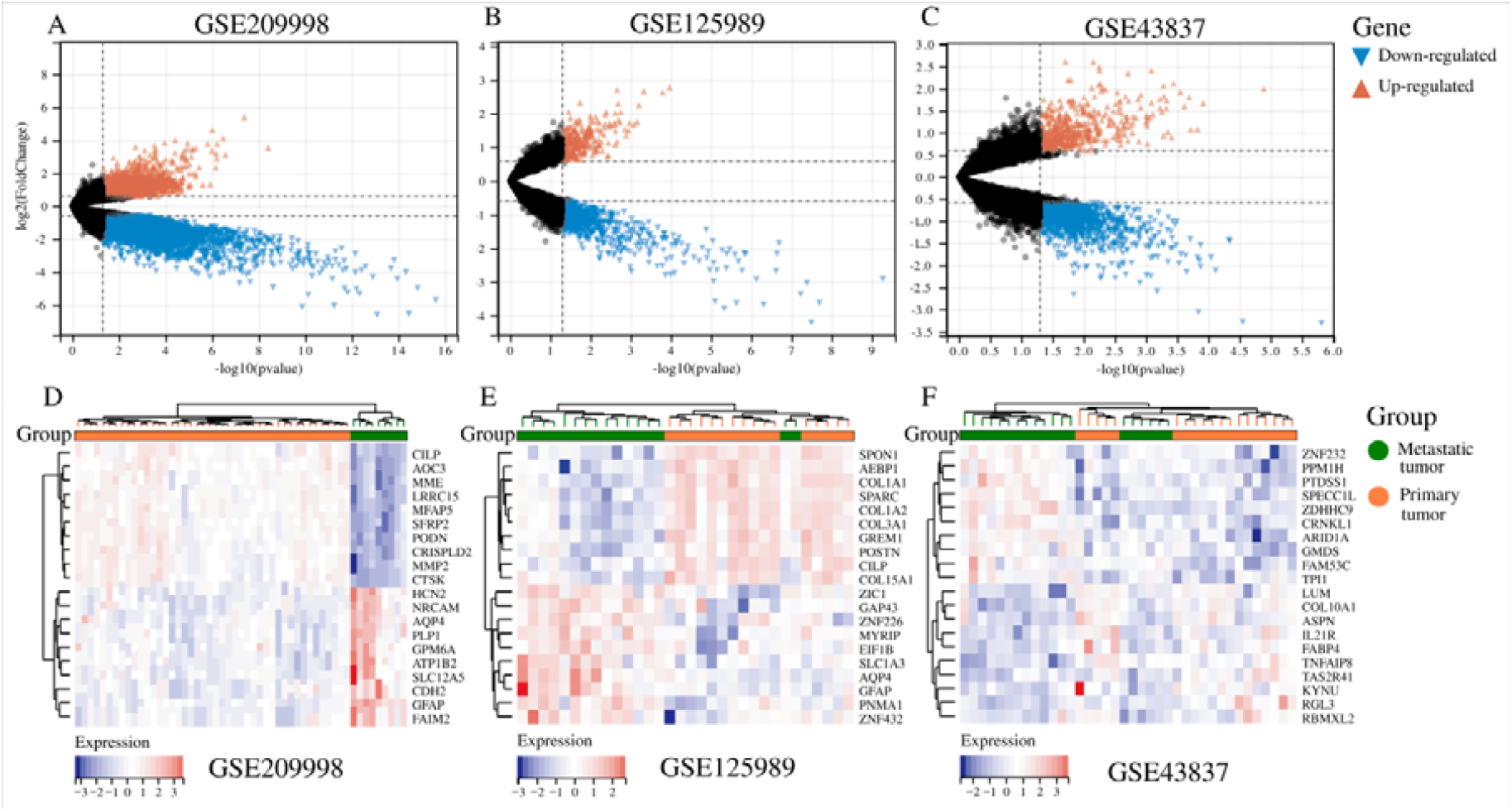
Identification of differentially expressed genes (DEGs) in three gene expression omnibus (GEO) datasets. (A-C) The volcano plots of differentially expressed genes in three datasets. Red represents up-regulated differential genes, black represents no significant difference genes, and blue represents down-regulated differential genes.(D-F)Heatmap of the top 10 upregulated and downregulated differentially expressed genes in three datasets. Blue represents down-regulated expression and red up-regulated expression. The green color of the top bars represents the metastatic tumor group, and the orange color of the top bars represents the primary tumor group.

Heatmaps were generated for the top 10 upregulated genes (GFAP, AQP4, COX6B1, TUBB2B, GPM6A, PPM1H,SLC6A1, AARS, HCN2, SPECC1L) and top 10 downregulated genes (LUM, MFAP5, FABP4, COL10A1, SERPINF1, CILP, SULF1, COL6A3, DPT, CTSK) (**Figure 3**).

**Figure 3.**
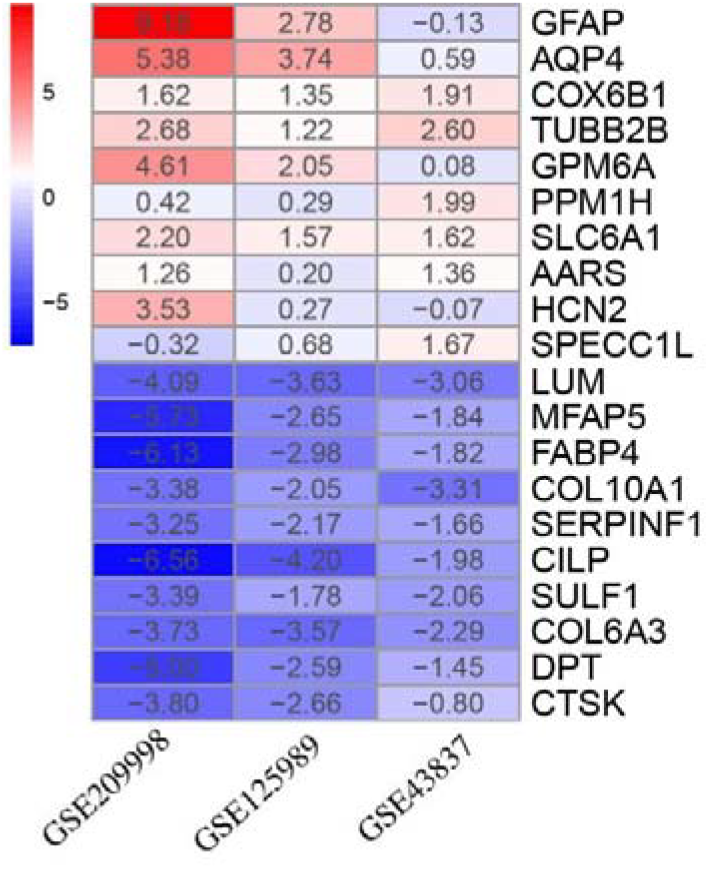
Heatmap shows the top 10 commonly upregulated and top 10 commonly downregulated genes in the RRA analysis of 3 bulk tissue mRNA-seq datasets. Red represents up-regulated genes, while blue represents down-regulated genes. The depth of the color represents the size of the log2 FC.

### Pathway analysis of commonly upregulated/downregulated genes

According to the cutoff criteria, pathway analysis was performed on the top 10 commonly DEGs in terms of upregulation and downregulation. The GO analysis results showed that the commonly differentially expressed genes were significantly enriched in pathways related to the extracellular region, extracellular matrix, and extracellular structural organization. The genes enriched in these pathways were downregulated in the metastatic foci compared to the primary foci (**Figure 4A, Supplementary Table 1**). Hallmark pathway analysis revealed that LUM, MFAP5, and COL6A3 were enriched in the epithelial-mesenchymal transition pathway (adj P < 0.05) (**Figure 4B, Supplementary Table 2**). These three genes were downregulated in the metastatic foci compared to the primary foci. No significant enrichment of meaningful pathways was found in the KEGG analysis (**Supplementary Table 3**).

**Figure 4.**
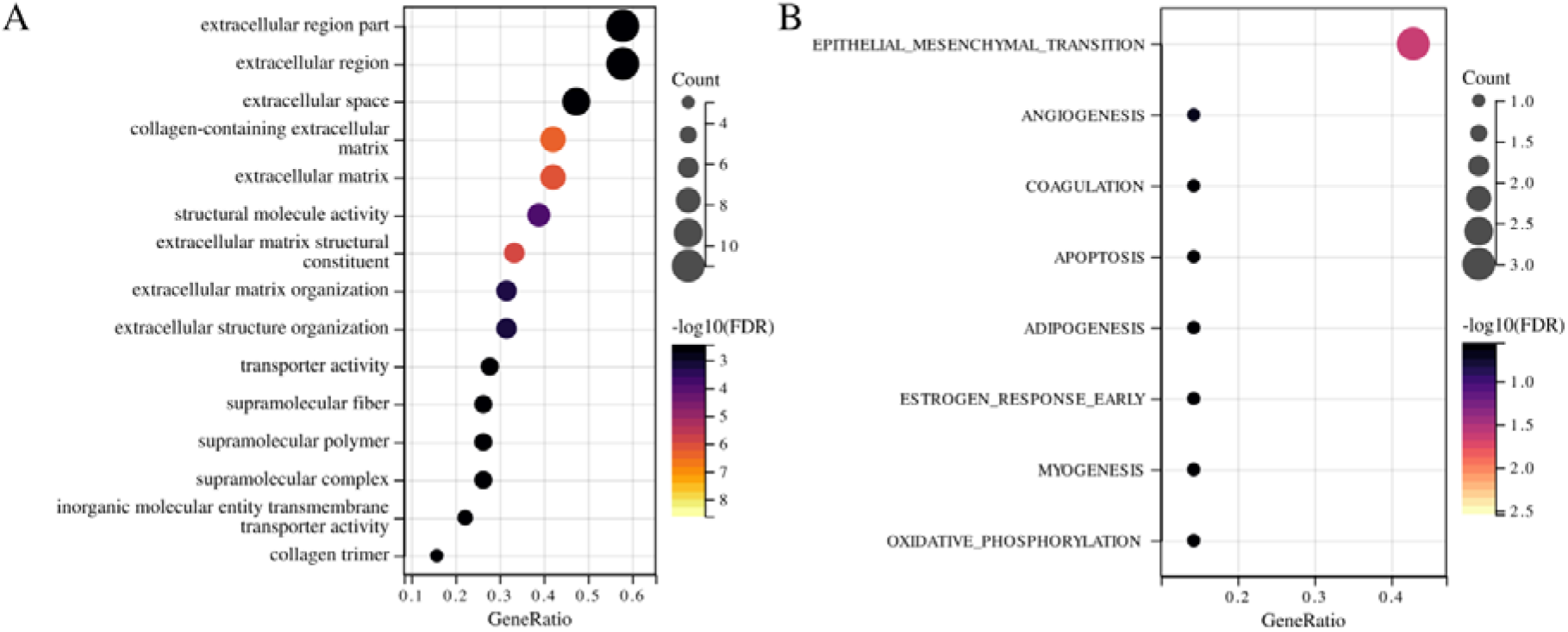
GO and Hallmark pathway analysis of commonly upregulated/downregulated genes. (A) GO pathway analysis. (B) Hallmark pathway analysis.

### scRNA-seq clustering analysis revealed heterogeneous cell populations in animal models of human BCBM

According to the visualization of P-value distributions for the 20 principal components using the JackStraw function, a total of 14 important principal components were determined and retained for visualization and clustering (**Figure 5**). All cells were projected onto a two-dimensional plane using the UMAP method, and further identified into 9 cell clusters based on cell types (**Figure 6A**). By considering the tissue origin, it can be observed that dormant BC cells in the tumor microenvironment are mainly concentrated in cluster 6, while the majority of proliferating BC cells are composed of the remaining clusters (**Figure 6B, C**). The stacked bar plot of cell proportions in different groups reveals that both the proliferating metastatic tumor cell population and dormant metastatic tumor cell population consist of distinct heterogeneous BC cell clusters, and there are significant differences in the composition proportions of cell clusters. Subcluster 6 dominates in the dormant tumor cell population, while clusters 0, 1, 2, 3, 4, and 5 dominate in the proliferating tumor cell population. Clusters 7 and 8 are present in both the proliferating tumor cell population and dormant tumor cell population, occupying a certain proportion in each (**Figure 6D**).

**Figure 5.**
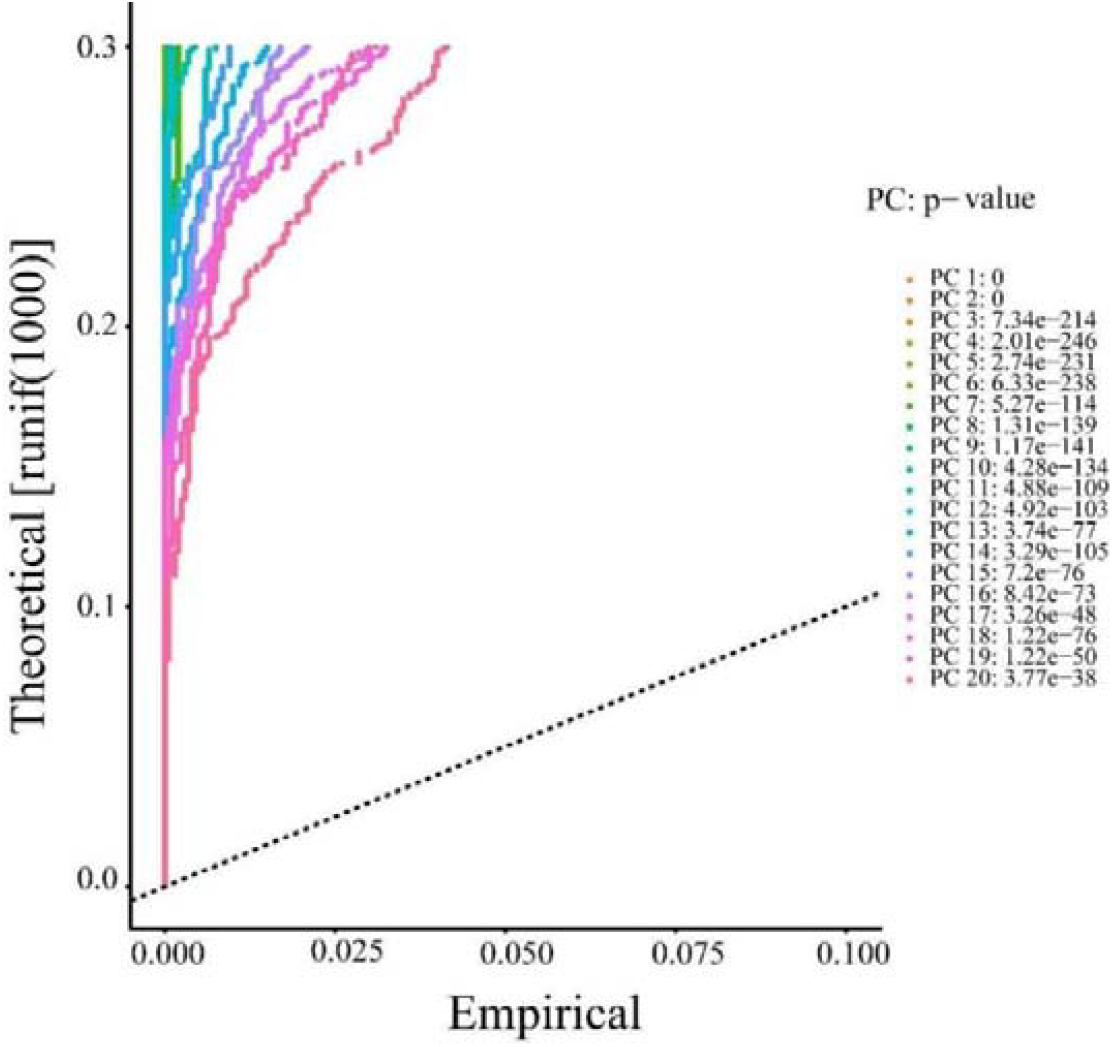
Jackstraw plot shows the p-value distribution of each principal component (PC). Significant PCs were defined as those above the dashed line with statistically significant p-values (P < 0.05).

**Figure 6.**
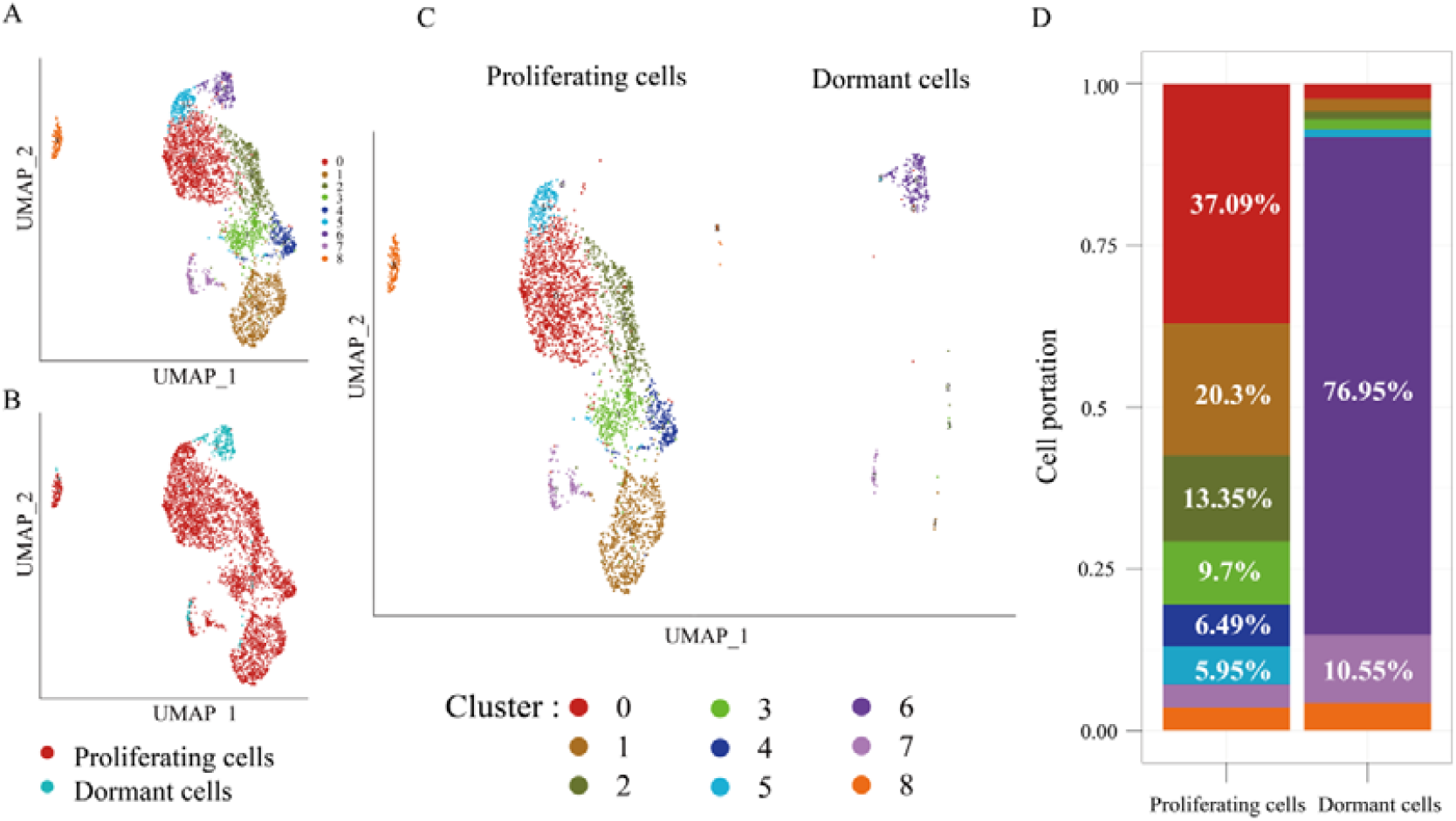
scRNA-seq analysis reveals cell cluster heterogeneity in animal models of human BCBM. (A) Cells were clustered into 9 types by the UMAP dimensionality reduction algorithm, with each color representing a cluster. (B) Cells were labeled with two colors according to their source. Red represents proliferating cells, and blue represents dormant cells. (C-D) Cells were labeled with nine colors according to their clusters. Bar charts show the proportion of cell clusters in proliferating cells and dormant cells.

### GSVA analysis based on scRNA-seq data revealed metabolic pathway differences between clusters of proliferating tumor cells and dormant tumor cells

Pathway analysis revealed that compared to the primary tumor, the metastatic lesion exhibited significant weakening of pathways related to ECM and protein digestion and absorption. When comparing proliferating tumor cells to dormant tumor cells, the pathways associated with ECM and structure, apoptosis, and inflammatory response were significantly attenuated. However, pathways involved in cell cycle regulation, tumor cell proliferation, and hypoxia response were significantly enhanced in proliferating tumor cells compared to dormant metastatic cells (**Figure 7A, Supplementary Table 4**). Further analysis was performed on different heterogeneous subpopulations of metastatic BC cells. The results showed that Subcluster 6 was a unique subpopulation of dormant tumor cells, exhibiting enhanced functions in EMT-related genes, ECM-related genes, collagen formation, tumor inflammation index (TIS), and IL-10 anti-inflammatory signaling pathway. Among the heterogeneous cancer cell subpopulations constituting the proliferating tumor metastatic group, cluster 3 showed significant enhancement in DNA replication pathway, while cluster 4 exhibited significant enhancement in tumor proliferation signaling pathway, and cluster 4 was uniquely associated with proliferating tumor cells (**Figure 7B, Supplementary Table 5**).

**Figure 7.**
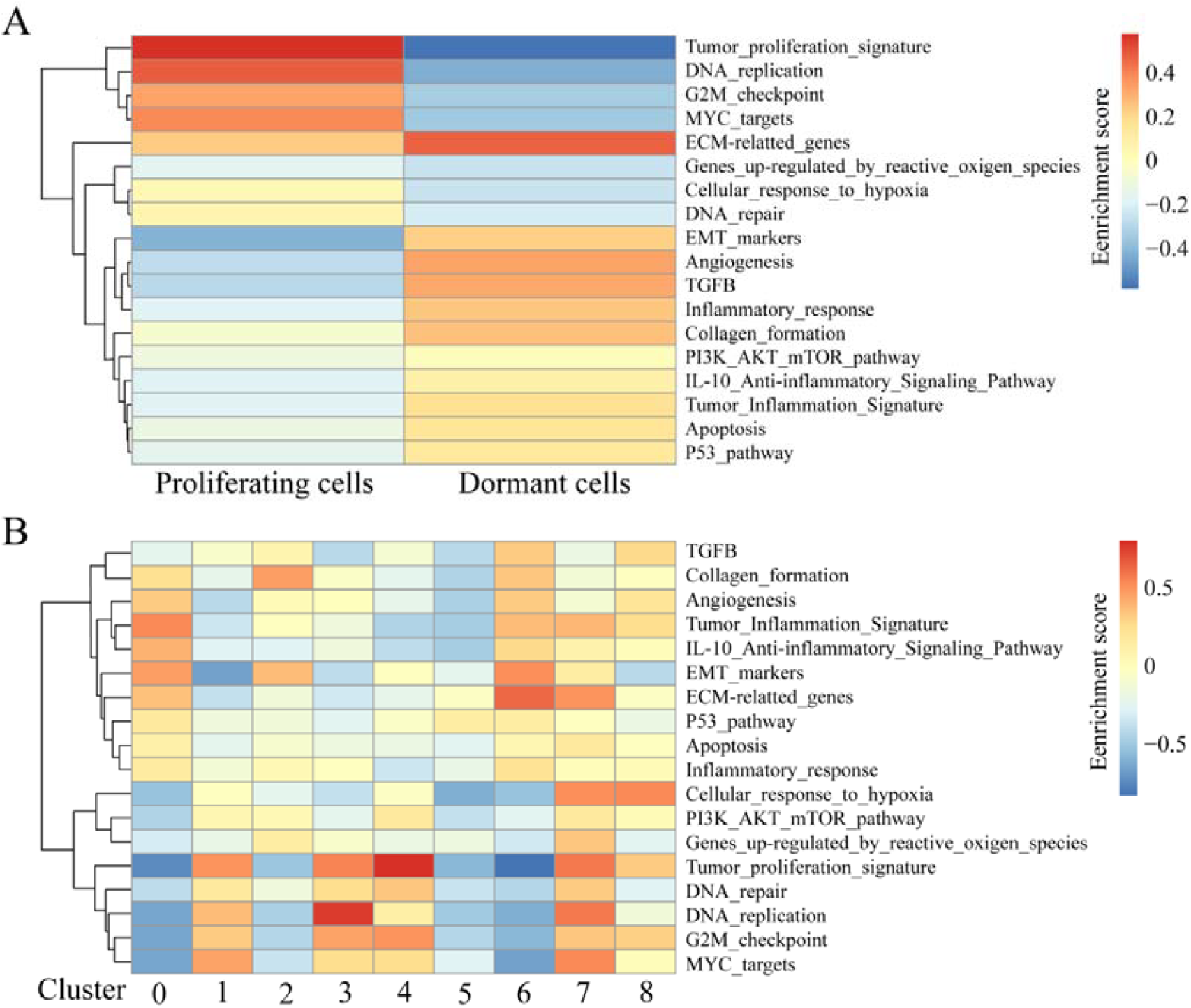
The heatmap displays the results of GSVA enrichment analysis performed on the scRNA-seq data. Red color represents positive enrichment scores (pathway activation), while blue color represents negative enrichment scores (pathway inhibition). The depth of color represents the magnitude of the enrichment score, with darker colors indicating higher scores and lighter colors indicating lower scores. (A) Divide the heatmap into two columns based on cell sources (proliferating cells and dormant cells). Each row represents a metabolic pathway. (B) Divide the heatmap into nine columns based on cell clusters (0-8). Each row represents a metabolic pathway.

## Discussion

BC is the most common cancer among women worldwide. According to previous research statistics, approximately 10-30% of patients with metastatic BC have brain metastases (16, 17). The process of metastasis involves a complex series of steps, including invasion, survival, and proliferation of cancer cells in distant organs. Recent studies have emphasized the importance of the tumor microenvironment (TME) in this process. The TME of brain metastases is a complex network composed of cells, signaling molecules, and ECM components that support the growth and survival of cancer cells (18). The ECM is a major component of the tumor microenvironment, providing structural support and regulating cell behavior (19). Changes in the structure and composition of the TME, particularly ECM remodeling, are closely associated with cancer growth, progression, and metastasis (20, 21). In BC, ECM plays a crucial role in invasion and metastasis. Some studies have found that ECM degradation is a common feature of BC invasion and metastasis (22), while others suggest that ECM stiffness and enhancement promote cancer cell migration and invasion (23, 24). The reason for this contradiction may be the different remodeling of the tumor microenvironment by BC cells at different stages of metastasis, exhibiting different biological behaviors.

Our research found that compared to the primary tumor, the metastatic lesion exhibited significant functional pathway attenuation in ECM, extracellular structure, and protein digestion and absorption. We believe that the ECM stiffening caused by BC cells before metastasis may promote cancer cell metastasis. Previous studies have shown that ECM stiffening, characterized by increased collagen deposition and thickening of interstitial collagens, can enhance the invasiveness of breast cells (23, 25). However, during the process of cancer cell colonization and progression in the brain, excessive ECM may hinder cancer cell dissemination. Our research discovered that brain metastatic cancer cells downregulated the extracellular region of ECM, extracellular structure-related pathways, and EMT pathway. Therefore, ECM degradation may be one of the adaptive behaviors of BCBM to the brain microenvironment, favoring the development of metastatic lesions. Single-cell analysis also confirmed the downregulation of ECM and EMT in the dormant cancer cell population of metastatic lesions, while dormant cancer cells did not proliferate upon brain colonization and retained the biological characteristics of the primary tumor, namely ECM stiffening capability. These facts support the notion that BC cells exhibit different remodeling behaviors of the tumor microenvironment at different stages of metastasis.

Once BC cells establish themselves in the brain, they must adapt to the unique characteristics of the brain TME. In a study by Valiente et al., brain metastatic cells exhibited specific phenotypic changes, including increased proliferation, decreased apoptosis, and enhanced hypoxia response (26). These changes may be related to the brain TME, which is characterized by a low-oxygen environment. Enhancing anti-hypoxia pathways may also be a means by which BC cells adapt to the environment. Another study by Dekker et al. found that the YAP/TAZ mechanotransduction pathway serves as an important link between hypoxia and ECM in the tumor microenvironment, collectively driving BC progression (27). This study indirectly supports our findings.

The heterogeneity of cancer cells within tumors can lead to different functions and behaviors of tumor cells. As early as 1984, Heppne’s research indicated that cancer cells within tumors can have different phenotypes and functions, including the ability to invade, grow, and metastasize (28). We analyzed cell subgroups and the results showed that cluster 3 exhibited significantly enhanced DNA replication pathways, while cluster 4 showed significantly enhanced tumor proliferation signaling pathways. Cluster 4 was identified as a unique subgroup of proliferating metastatic cells, similar to the highly proliferative characteristics of proliferating metastatic cancer cells. cluster 1, 3, 4, and 5 exhibited downregulation of ECM construction abilities such as key genes involved in EMT, ECM-related genes, and collagen formation pathways compared to dormant metastatic lesions. Furthermore, their tumor inflammation pathway functions were significantly weakened compared to other subgroups. These behaviors may facilitate tumor cell migration and immune escape. Cluster 6 was identified as a unique subgroup of dormant metastatic lesion cancer cells, which exhibited upregulation of ECM construction abilities and significantly enhanced tumor inflammation pathway function compared to other subgroups. The enhanced matrix and inflammation may cause difficulties in tumor cell migration and aggregation of surrounding immune cells, which could be the reason for these cancer cells entering a dormant state.

Zou et al.’s research suggests that the metastatic ecosystem formed during BCBM exhibits significant immune suppressive cell reprogramming to aid immune escape of brain metastatic cells (29). Another study found the absence of cellular immunity in the microenvironment of BCBM, which may be an important condition for cancer cells to establish themselves in the brain (30). We believe that primary tumor cells may consist of different heterogeneous cancer cell populations, and upon entering the brain environment, populations that adapt to the environment can survive and develop, while those that are not adaptable may gradually become dormant or even undergo apoptosis. proliferating BC metastatic cells, by reducing inflammatory responses for immune escape, facilitate the growth and colonization of metastatic cancer cells in the brain TME.

In summary, the process of BC metastasis to the brain is complex. This study found that the characteristics of cancer cells at different stages and the heterogeneity of different subgroups of cancer cells play important roles in the process of BC cell colonization and growth from the primary site to brain metastatic lesions. This effect can be achieved through adaptation to the brain environment and reshaping of the tumor microenvironment. By gaining a deeper understanding of these underlying mechanisms, researchers and clinicians can develop more effective treatment strategies, ultimately improving the prognosis of BCBM patients.

## Supporting information

GO enrichment analysis results of DEGs in bulk tissue sequencing data.

Hallmark enrichment analysis results of DEGs in bulk tissue sequencing data.

KEGG enrichment analysis results of DEGs in bulk tissue sequencing data.

Results of GSVA enrichment analysis performed on the scRNA-seq data, which were grouped based on cell sources (proliferating cells and dormant cells).

Results of GSVA enrichment analysis performed on the scRNA-seq data, which were grouped based on cell cluster.

## Supplementary Materials

Supplementary Table 1. GO enrichment analysis results of DEGs in bulk tissue sequencing data.

Supplementary Table 2. Hallmark enrichment analysis results of DEGs in bulk tissue sequencing data.

Supplementary Table 3. KEGG enrichment analysis results of DEGs in bulk tissue sequencing data.

Supplementary Table 4. Results of GSVA enrichment analysis performed on the scRNA-seq data, which were grouped based on cell sources (proliferating cells and dormant cells).

Supplementary Table 5. Results of GSVA enrichment analysis performed on the scRNA-seq data, which were grouped based on cell cluster.

## Acknowledgments

We thank GEO for generating the scRNA-seq data and mRNA-seq data.

## Author Contributions

ZY and ZYH wrote the manuscript, designed the study and interpreted the data. XW performed the statistical analysis and interpreted the results. XW and RL was involved in designing the study and critically revising the manuscript. All authors read and approved the final manuscript.

## Ethics approval and consent to participate

Not applicable.

## Availability of data and materials

The datasets analyzed during the current study are available in Gene Expression Omnibus (GEO, https://www.ncbi.nlm.nih.gov/geo/).

