## Supplementary material for "Differential Adaptation and Remodeling of the Tumor Microenvironment in Breast Cancer Cells During the Stages of Brain Metastasis": GO enrichment analysis results of DEGs in bulk tissue sequencing data.

### Supplementary Table 1. GO enrichment analysis results of DEGs in bulk tissue sequencing data.

| ID | Pathway | GeneRatio | BgRatio | pvalue | p.adjust | qvalue | geneID | Count | ONT |
| --- | --- | --- | --- | --- | --- | --- | --- | --- | --- |
| GO:0062023 | collagen-containing extracellular matrix | 8/19 | 399/18675 | 2.49115381567067e-09 | 3.6869076471926e-07 | 2.59604450264628e-07 | LUM/MFAP5/COL10A1/SERPINF1/CILP/SULF1/COL6A3/DPT | 8 | CC |
| GO:0031012 | extracellular matrix | 8/19 | 468/18675 | 8.69257518177506e-09 | 6.43250563451355e-07 | 4.52928917366175e-07 | LUM/MFAP5/COL10A1/SERPINF1/CILP/SULF1/COL6A3/DPT | 8 | CC |
| GO:0005201 | extracellular matrix structural constituent | 6/18 | 158/16967 | 1.00343223324784e-08 | 1.07367248957518e-06 | 7.07683785553737e-07 | LUM/MFAP5/COL10A1/CILP/COL6A3/DPT | 6 | MF |
| GO:0005198 | structural molecule activity | 7/18 | 653/16967 | 2.66023890179152e-06 | 0.000142322781245847 | 9.38084244315958e-05 | GFAP/LUM/MFAP5/COL10A1/CILP/COL6A3/DPT | 7 | MF |
| GO:0030198 | extracellular matrix organization | 6/19 | 334/17910 | 8.89068725288633e-07 | 0.000671246887592918 | 0.000550286747862859 | LUM/MFAP5/COL10A1/SULF1/COL6A3/CTSK | 6 | BP |
| GO:0043062 | extracellular structure organization | 6/19 | 387/17910 | 2.09367473563201e-06 | 0.000790362212701082 | 0.000647937233974537 | LUM/MFAP5/COL10A1/SULF1/COL6A3/CTSK | 6 | BP |
| GO:0005581 | collagen trimer | 3/19 | 86/18675 | 8.66233880908265e-05 | 0.00368526656518171 | 0.00259488897548356 | LUM/COL10A1/COL6A3 | 3 | CC |
| GO:0044421 | extracellular region part | 11/19 | 3312/18675 | 9.96017990589651e-05 | 0.00368526656518171 | 0.00259488897548356 | GPM6A/LUM/MFAP5/FABP4/COL10A1/SERPINF1/CILP/SULF1/COL6A3/DPT/CTSK | 11 | CC |
| GO:0044420 | extracellular matrix component | 2/19 | 47/18675 | 0.00103156862863166 | 0.0218752478838201 | 0.0154029128058193 | LUM/MFAP5 | 2 | CC |
| GO:0005576 | extracellular region | 11/19 | 4295/18675 | 0.00109244259410598 | 0.0218752478838201 | 0.0154029128058193 | GPM6A/LUM/MFAP5/FABP4/COL10A1/SERPINF1/CILP/SULF1/COL6A3/DPT/CTSK | 11 | CC |
| GO:0099512 | supramolecular fiber | 5/19 | 845/18675 | 0.00128209544771797 | 0.0218752478838201 | 0.0154029128058193 | GFAP/TUBB2B/LUM/MFAP5/COL6A3 | 5 | CC |
| GO:0099081 | supramolecular polymer | 5/19 | 851/18675 | 0.00132329063833896 | 0.0218752478838201 | 0.0154029128058193 | GFAP/TUBB2B/LUM/MFAP5/COL6A3 | 5 | CC |
| GO:0099080 | supramolecular complex | 5/19 | 852/18675 | 0.00133025156050258 | 0.0218752478838201 | 0.0154029128058193 | GFAP/TUBB2B/LUM/MFAP5/COL6A3 | 5 | CC |
| GO:0005615 | extracellular space | 9/19 | 3051/18675 | 0.00157449724211684 | 0.0233025591833292 | 0.0164079186283755 | GPM6A/LUM/FABP4/SERPINF1/CILP/SULF1/COL6A3/DPT/CTSK | 9 | CC |
| GO:0030020 | extracellular matrix structural constituent conferring tensile strength | 2/18 | 41/16967 | 0.000850560226337338 | 0.0303366480726984 | 0.0199956263735444 | COL10A1/COL6A3 | 2 | MF |
| GO:0099056 | integral component of presynaptic membrane | 2/19 | 74/18675 | 0.00253567131510704 | 0.0313497689323773 | 0.0220741616237934 | GPM6A/SLC6A1 | 2 | CC |
| GO:0005589 | collagen type VI trimer | 1/19 | 3/18675 | 0.0030492676825864 | 0.0313497689323773 | 0.0220741616237934 | COL6A3 | 1 | CC |
| GO:0098647 | collagen beaded filament | 1/19 | 3/18675 | 0.0030492676825864 | 0.0313497689323773 | 0.0220741616237934 | COL6A3 | 1 | CC |
| GO:0098855 | HCN channel complex | 1/19 | 3/18675 | 0.0030492676825864 | 0.0313497689323773 | 0.0220741616237934 | HCN2 | 1 | CC |
| GO:0098889 | intrinsic component of presynaptic membrane | 2/19 | 83/18675 | 0.00317734144584905 | 0.0313497689323773 | 0.0220741616237934 | GPM6A/SLC6A1 | 2 | CC |
| GO:0043202 | lysosomal lumen | 2/19 | 94/18675 | 0.00405407259760963 | 0.0375001715278891 | 0.0264048149449575 | LUM/CTSK | 2 | CC |
| GO:0036021 | endolysosome lumen | 1/19 | 5/18675 | 0.00507721683134765 | 0.0417460050577474 | 0.029394413234118 | CTSK | 1 | CC |
| GO:0097450 | astrocyte end-foot | 1/19 | 5/18675 | 0.00507721683134765 | 0.0417460050577474 | 0.029394413234118 | AQP4 | 1 | CC |
| GO:0005518 | collagen binding | 2/18 | 61/16967 | 0.00187451383153867 | 0.044365524930905 | 0.0292424020695587 | LUM/CTSK | 2 | MF |
| GO:0005215 | transporter activity | 5/18 | 908/16967 | 0.00207315537060304 | 0.044365524930905 | 0.0292424020695587 | AQP4/COX6B1/GPM6A/HCN2/FABP4 | 5 | MF |
| GO:0005222 | intracellular cAMP-activated cation channel activity | 1/18 | 3/16967 | 0.00317946064676877 | 0.0477294459121358 | 0.0314596446248214 | HCN2 | 1 | MF |
| GO:0008449 | N-acetylglucosamine-6-sulfatase activity | 1/18 | 3/16967 | 0.00317946064676877 | 0.0477294459121358 | 0.0314596446248214 | SULF1 | 1 | MF |
| GO:0015318 | inorganic molecular entity transmembrane transporter activity | 4/18 | 619/16967 | 0.00356855670371109 | 0.0477294459121358 | 0.0314596446248214 | AQP4/COX6B1/GPM6A/HCN2 | 4 | MF |
| GO:0042383 | sarcolemma | 2/19 | 125/18675 | 0.00705457585395906 | 0.0549514329676811 | 0.0386926875092491 | AQP4/COL6A3 | 2 | CC |
| GO:0022838 | substrate-specific channel activity | 3/18 | 338/16967 | 0.00512032895116282 | 0.0566435999034509 | 0.0373351814415269 | AQP4/GPM6A/HCN2 | 3 | MF |
| GO:0005324 | long-chain fatty acid transporter activity | 1/18 | 5/16967 | 0.00529379438350008 | 0.0566435999034509 | 0.0373351814415269 | FABP4 | 1 | MF |
| GO:0015267 | channel activity | 3/18 | 370/16967 | 0.00658061291854321 | 0.0591162399526348 | 0.0389649589456619 | AQP4/GPM6A/HCN2 | 3 | MF |
| GO:0022803 | passive transmembrane transporter activity | 3/18 | 371/16967 | 0.0066298586862768 | 0.0591162399526348 | 0.0389649589456619 | AQP4/GPM6A/HCN2 | 3 | MF |
| GO:0043203 | axon hillock | 1/19 | 8/18675 | 0.00811181441318476 | 0.0600274266575672 | 0.0422668224686995 | SERPINF1 | 1 | CC |
| GO:0042734 | presynaptic membrane | 2/19 | 141/18675 | 0.00889791131486098 | 0.0627090892666393 | 0.0441550486301372 | GPM6A/SLC6A1 | 2 | CC |
| GO:0005221 | intracellular cyclic nucleotide activated cation channel activity | 1/18 | 8/16967 | 0.00845735495350186 | 0.0644448496176901 | 0.0424771758424519 | HCN2 | 1 | MF |
| GO:0043855 | cyclic nucleotide-gated ion channel activity | 1/18 | 8/16967 | 0.00845735495350186 | 0.0644448496176901 | 0.0424771758424519 | HCN2 | 1 | MF |
| GO:0015250 | water channel activity | 1/18 | 9/16967 | 0.00950976179416707 | 0.0644448496176901 | 0.0424771758424519 | AQP4 | 1 | MF |
| GO:0022857 | transmembrane transporter activity | 4/18 | 838/16967 | 0.0103781437837907 | 0.0644448496176901 | 0.0424771758424519 | AQP4/COX6B1/GPM6A/HCN2 | 4 | MF |
| GO:0005372 | water transmembrane transporter activity | 1/18 | 10/16967 | 0.0105611136214876 | 0.0644448496176901 | 0.0424771758424519 | AQP4 | 1 | MF |
| GO:0022890 | inorganic cation transmembrane transporter activity | 3/18 | 444/16967 | 0.0108411896553123 | 0.0644448496176901 | 0.0424771758424519 | COX6B1/GPM6A/HCN2 | 3 | MF |
| GO:0001527 | microfibril | 1/19 | 10/18675 | 0.0101300057451985 | 0.0659348403846668 | 0.0464263812096872 | MFAP5 | 1 | CC |
| GO:0099699 | integral component of synaptic membrane | 2/19 | 152/18675 | 0.0102772435691404 | 0.0659348403846668 | 0.0464263812096872 | GPM6A/SLC6A1 | 2 | CC |
| GO:0005583 | fibrillar collagen trimer | 1/19 | 11/18675 | 0.0111376419568694 | 0.0659348403846668 | 0.0464263812096872 | LUM | 1 | CC |
| GO:0098643 | banded collagen fibril | 1/19 | 11/18675 | 0.0111376419568694 | 0.0659348403846668 | 0.0464263812096872 | LUM | 1 | CC |
| GO:0099240 | intrinsic component of synaptic membrane | 2/19 | 164/18675 | 0.0118834327891594 | 0.066568790530203 | 0.0468727614686351 | GPM6A/SLC6A1 | 2 | CC |
| GO:0097449 | astrocyte projection | 1/19 | 12/18675 | 0.01214430638051 | 0.066568790530203 | 0.0468727614686351 | AQP4 | 1 | CC |
| GO:0005775 | vacuolar lumen | 2/19 | 171/18675 | 0.0128683011067856 | 0.0680181629930096 | 0.0478933011117208 | LUM/CTSK | 2 | CC |
| GO:0004065 | arylsulfatase activity | 1/18 | 12/16967 | 0.012660656217103 | 0.0712994850121063 | 0.046995233603651 | SULF1 | 1 | MF |
| GO:0099059 | integral component of presynaptic active zone membrane | 1/19 | 14/18675 | 0.0141547234037187 | 0.0722378987500127 | 0.0508645233019293 | GPM6A | 1 | CC |
| GO:0008324 | cation transmembrane transporter activity | 3/18 | 484/16967 | 0.0136830741741479 | 0.0732044468316913 | 0.0482508405088373 | COX6B1/GPM6A/HCN2 | 3 | MF |
| GO:0036041 | long-chain fatty acid binding | 1/18 | 14/16967 | 0.0147559906935575 | 0.0751852859147929 | 0.049556459973351 | FABP4 | 1 | MF |
| GO:0070062 | extracellular exosome | 6/19 | 2162/18675 | 0.0170953698184108 | 0.0768074961127827 | 0.054082091857507 | GPM6A/LUM/FABP4/SERPINF1/CILP/COL6A3 | 6 | CC |
| GO:1903561 | extracellular vesicle | 6/19 | 2185/18675 | 0.0179509826473051 | 0.0768074961127827 | 0.054082091857507 | GPM6A/LUM/FABP4/SERPINF1/CILP/COL6A3 | 6 | CC |
| GO:0043230 | extracellular organelle | 6/19 | 2187/18675 | 0.0180267976198261 | 0.0768074961127827 | 0.054082091857507 | GPM6A/LUM/FABP4/SERPINF1/CILP/COL6A3 | 6 | CC |
| GO:0036019 | endolysosome | 1/19 | 18/18675 | 0.0181639348915364 | 0.0768074961127827 | 0.054082091857507 | CTSK | 1 | CC |
| GO:0098644 | complex of collagen trimers | 1/19 | 18/18675 | 0.0181639348915364 | 0.0768074961127827 | 0.054082091857507 | LUM | 1 | CC |
| GO:0098945 | intrinsic component of presynaptic active zone membrane | 1/19 | 18/18675 | 0.0181639348915364 | 0.0768074961127827 | 0.054082091857507 | GPM6A | 1 | CC |
| GO:0008484 | sulfuric ester hydrolase activity | 1/18 | 15/16967 | 0.0158020823686301 | 0.0768555824292465 | 0.0506573932391492 | SULF1 | 1 | MF |
| GO:0005248 | voltage-gated sodium channel activity | 1/18 | 17/16967 | 0.0178911195478486 | 0.0803142945008543 | 0.0529371149194023 | HCN2 | 1 | MF |
| GO:0004129 | cytochrome-c oxidase activity | 1/18 | 19/16967 | 0.0199759684294397 | 0.0803142945008543 | 0.0529371149194023 | COX6B1 | 1 | MF |
| GO:0015002 | heme-copper terminal oxidase activity | 1/18 | 19/16967 | 0.0199759684294397 | 0.0803142945008543 | 0.0529371149194023 | COX6B1 | 1 | MF |
| GO:0016676 | oxidoreductase activity, acting on a heme group of donors, oxygen as acceptor | 1/18 | 19/16967 | 0.0199759684294397 | 0.0803142945008543 | 0.0529371149194023 | COX6B1 | 1 | MF |
| GO:0016675 | oxidoreductase activity, acting on a heme group of donors | 1/18 | 20/16967 | 0.0210168247291955 | 0.0803142945008543 | 0.0529371149194023 | COX6B1 | 1 | MF |
| GO:0030552 | cAMP binding | 1/18 | 20/16967 | 0.0210168247291955 | 0.0803142945008543 | 0.0529371149194023 | HCN2 | 1 | MF |
| GO:0030021 | extracellular matrix structural constituent conferring compression resistance | 1/18 | 21/16967 | 0.0220566369174811 | 0.081381384488637 | 0.0536404600171046 | LUM | 1 | MF |
| GO:0001968 | fibronectin binding | 1/18 | 24/16967 | 0.0251698186673962 | 0.0897723532470465 | 0.0591711526566858 | CTSK | 1 | MF |
| GO:0005217 | intracellular ligand-gated ion channel activity | 1/18 | 25/16967 | 0.0262054642593215 | 0.0904511185724967 | 0.0596185434762152 | HCN2 | 1 | MF |
| GO:0045202 | synapse | 4/19 | 1091/18675 | 0.0221988776752952 | 0.0912620526651024 | 0.064259909060065 | TUBB2B/GPM6A/PPM1H/SLC6A1 | 4 | CC |
| GO:0048787 | presynaptic active zone membrane | 1/19 | 23/18675 | 0.0231537296544618 | 0.0926149186178473 | 0.0652124960395938 | GPM6A | 1 | CC |
| GO:0097386 | glial cell projection | 1/19 | 24/18675 | 0.024148801450332 | 0.0940532267012932 | 0.0662252449745948 | AQP4 | 1 | CC |
| GO:0005261 | cation channel activity | 2/18 | 254/16967 | 0.0291703844999183 | 0.0950230354565336 | 0.0626320056624471 | GPM6A/HCN2 | 2 | MF |
| GO:0005504 | fatty acid binding | 1/18 | 28/16967 | 0.0293061698136973 | 0.0950230354565336 | 0.0626320056624471 | FABP4 | 1 | MF |
| GO:0015077 | monovalent inorganic cation transmembrane transporter activity | 2/18 | 266/16967 | 0.0317591746397695 | 0.0990126561031262 | 0.0652616621633985 | COX6B1/HCN2 | 2 | MF |
| GO:0015075 | ion transmembrane transporter activity | 3/18 | 672/16967 | 0.0323873174169104 | 0.0990126561031262 | 0.0652616621633985 | COX6B1/GPM6A/HCN2 | 3 | MF |
| GO:0005887 | integral component of plasma membrane | 4/19 | 1158/18675 | 0.0269776067626774 | 0.0990877407219271 | 0.0697701730545575 | AQP4/GPM6A/SLC6A1/HCN2 | 4 | CC |
| GO:0000323 | lytic vacuole | 3/19 | 659/18675 | 0.0278159536215677 | 0.0990877407219271 | 0.0697701730545575 | GFAP/LUM/CTSK | 3 | CC |
| GO:0005764 | lysosome | 3/19 | 659/18675 | 0.0278159536215677 | 0.0990877407219271 | 0.0697701730545575 | GFAP/LUM/CTSK | 3 | CC |
| GO:0044295 | axonal growth cone | 1/19 | 28/18675 | 0.028119493988655 | 0.0990877407219271 | 0.0697701730545575 | GPM6A | 1 | CC |
| GO:0030551 | cyclic nucleotide binding | 1/18 | 32/16967 | 0.0334259409543508 | 0.0993493245032094 | 0.0654835685363013 | HCN2 | 1 | MF |
