## Supplementary material for "Differential Adaptation and Remodeling of the Tumor Microenvironment in Breast Cancer Cells During the Stages of Brain Metastasis": Hallmark enrichment analysis results of DEGs in bulk tissue sequencing data.

### Supplementary Table 2. Hallmark enrichment analysis results of DEGs in bulk tissue sequencing data.

| Description | GeneRatio | BgRatio | pvalue | p.adjust | qvalue | geneID | Count |
| --- | --- | --- | --- | --- | --- | --- | --- |
| EPITHELIAL_MESENCHYMAL_TRANSITION | 3/7 | 200/4383 | 0.00285880475689797 | 0.0228704380551838 | 0.0210648771560903 | LUM/MFAP5/COL6A3 | 3 |
| ANGIOGENESIS | 1/7 | 36/4383 | 0.0561348800392154 | 0.224539520156862 | 0.206812715933952 | LUM | 1 |
| COAGULATION | 1/7 | 138/4383 | 0.20076249612173 | 0.279033363336343 | 0.257004413599264 | CTSK | 1 |
| APOPTOSIS | 1/7 | 161/4383 | 0.230607622268114 | 0.279033363336343 | 0.257004413599264 | LUM | 1 |
| ADIPOGENESIS | 1/7 | 200/4383 | 0.279033363336343 | 0.279033363336343 | 0.257004413599264 | FABP4 | 1 |
| ESTROGEN_RESPONSE_EARLY | 1/7 | 200/4383 | 0.279033363336343 | 0.279033363336343 | 0.257004413599264 | TUBB2B | 1 |
| MYOGENESIS | 1/7 | 200/4383 | 0.279033363336343 | 0.279033363336343 | 0.257004413599264 | COL6A3 | 1 |
| OXIDATIVE_PHOSPHORYLATION | 1/7 | 200/4383 | 0.279033363336343 | 0.279033363336343 | 0.257004413599264 | COX6B1 | 1 |
