## Supplementary material for "Differential Adaptation and Remodeling of the Tumor Microenvironment in Breast Cancer Cells During the Stages of Brain Metastasis": KEGG enrichment analysis results of DEGs in bulk tissue sequencing data.

### Supplementary Table 3. KEGG enrichment analysis results of DEGs in bulk tissue sequencing data.

| ID | Description | GeneRatio | BgRatio | pvalue | p.adjust | qvalue | geneID | Count | one_type |
| --- | --- | --- | --- | --- | --- | --- | --- | --- | --- |
| hsa04974 | Protein digestion and absorption | 2/13 | 95/7914 | 0.0102045899121614 | 0.336751467101325 | 0.336751467101325 | COL10A1/COL6A3 | 2 | Organismal Systems |
| hsa04962 | Vasopressin-regulated water reabsorption | 1/13 | 44/7914 | 0.0699656968007636 | 0.338488972116741 | 0.338488972116741 | AQP4 | 1 | Organismal Systems |
| hsa04923 | Regulation of lipolysis in adipocytes | 1/13 | 55/7914 | 0.0867363550761457 | 0.338488972116741 | 0.338488972116741 | FABP4 | 1 | Organismal Systems |
| hsa04929 | GnRH secretion New | 1/13 | 64/7914 | 0.100249687323194 | 0.338488972116741 | 0.338488972116741 | HCN2 | 1 | Organismal Systems |
| hsa00970 | Aminoacyl-tRNA biosynthesis | 1/13 | 66/7914 | 0.1032274740064 | 0.338488972116741 | 0.338488972116741 | AARS | 1 | Genetic Information Processing |
| hsa04976 | Bile secretion | 1/13 | 72/7914 | 0.11210633945809 | 0.338488972116741 | 0.338488972116741 | AQP4 | 1 | Organismal Systems |
| hsa03320 | PPAR signaling pathway | 1/13 | 76/7914 | 0.117980425067018 | 0.338488972116741 | 0.338488972116741 | FABP4 | 1 | Organismal Systems |
| hsa04721 | Synaptic vesicle cycle | 1/13 | 78/7914 | 0.12090399636588 | 0.338488972116741 | 0.338488972116741 | SLC6A1 | 1 | Organismal Systems |
| hsa04260 | Cardiac muscle contraction | 1/13 | 86/7914 | 0.132509058117147 | 0.338488972116741 | 0.338488972116741 | COX6B1 | 1 | Organismal Systems |
| hsa04512 | ECM-receptor interaction | 1/13 | 88/7914 | 0.135388142751793 | 0.338488972116741 | 0.338488972116741 | COL6A3 | 1 | Environmental Information Processing |
| hsa04540 | Gap junction | 1/13 | 88/7914 | 0.135388142751793 | 0.338488972116741 | 0.338488972116741 | TUBB2B | 1 | Cellular Processes |
| hsa04727 | GABAergic synapse | 1/13 | 89/7914 | 0.136824375072803 | 0.338488972116741 | 0.338488972116741 | SLC6A1 | 1 | Organismal Systems |
| hsa05323 | Rheumatoid arthritis | 1/13 | 93/7914 | 0.142547310015091 | 0.338488972116741 | 0.338488972116741 | CTSK | 1 | Human Diseases |
| hsa04620 | Toll-like receptor signaling pathway | 1/13 | 104/7914 | 0.158105283345089 | 0.338488972116741 | 0.338488972116741 | CTSK | 1 | Organismal Systems |
| hsa04142 | Lysosome | 1/13 | 123/7914 | 0.184365830826913 | 0.338488972116741 | 0.338488972116741 | CTSK | 1 | Cellular Processes |
| hsa04380 | Osteoclast differentiation | 1/13 | 128/7914 | 0.191149698787406 | 0.338488972116741 | 0.338488972116741 | CTSK | 1 | Organismal Systems |
| hsa00190 | Oxidative phosphorylation | 1/13 | 133/7914 | 0.19788145022877 | 0.338488972116741 | 0.338488972116741 | COX6B1 | 1 | Metabolism |
| hsa04210 | Apoptosis | 1/13 | 136/7914 | 0.201895638038981 | 0.338488972116741 | 0.338488972116741 | CTSK | 1 | Cellular Processes |
| hsa05012 | Parkinson disease | 1/13 | 142/7914 | 0.209868439957483 | 0.338488972116741 | 0.338488972116741 | COX6B1 | 1 | Human Diseases |
| hsa04932 | Non-alcoholic fatty liver disease (NAFLD) | 1/13 | 149/7914 | 0.219077094513302 | 0.338488972116741 | 0.338488972116741 | COX6B1 | 1 | Human Diseases |
| hsa04145 | Phagosome | 1/13 | 152/7914 | 0.222993249428303 | 0.338488972116741 | 0.338488972116741 | TUBB2B | 1 | Cellular Processes |
| hsa04310 | Wnt signaling pathway | 1/13 | 160/7914 | 0.23334787832927 | 0.338488972116741 | 0.338488972116741 | SERPINF1 | 1 | Environmental Information Processing |
| hsa04630 | Jak-STAT signaling pathway | 1/13 | 162/7914 | 0.235916556323789 | 0.338488972116741 | 0.338488972116741 | GFAP | 1 | Environmental Information Processing |
| hsa05010 | Alzheimer disease | 1/13 | 171/7914 | 0.247377595399736 | 0.3394962968195 | 0.3394962968195 | COX6B1 | 1 | Human Diseases |
| hsa05016 | Huntington disease | 1/13 | 193/7914 | 0.274728728937408 | 0.3394962968195 | 0.3394962968195 | COX6B1 | 1 | Human Diseases |
| hsa04510 | Focal adhesion | 1/13 | 199/7914 | 0.282027231764925 | 0.3394962968195 | 0.3394962968195 | COL6A3 | 1 | Cellular Processes |
| hsa05130 | Pathogenic Escherichia coli infection | 1/13 | 201/7914 | 0.284444959883068 | 0.3394962968195 | 0.3394962968195 | TUBB2B | 1 | Human Diseases |
| hsa05205 | Proteoglycans in cancer | 1/13 | 204/7914 | 0.288057463968061 | 0.3394962968195 | 0.3394962968195 | LUM | 1 | Human Diseases |
| hsa04024 | cAMP signaling pathway | 1/13 | 216/7914 | 0.302339784091146 | 0.344041823276132 | 0.344041823276132 | HCN2 | 1 | Environmental Information Processing |
| hsa04714 | Thermogenesis | 1/13 | 231/7914 | 0.319820715510916 | 0.351802787062007 | 0.351802787062007 | COX6B1 | 1 | Organismal Systems |
| hsa05165 | Human papillomavirus infection | 1/13 | 330/7914 | 0.42542970738126 | 0.452876785276825 | 0.452876785276825 | COL6A3 | 1 | Human Diseases |
| hsa04151 | PI3K-Akt signaling pathway | 1/13 | 354/7914 | 0.448641470570437 | 0.462661516525764 | 0.462661516525764 | COL6A3 | 1 | Environmental Information Processing |
| hsa01100 | Metabolic pathways | 1/13 | 1439/7914 | 0.92654589625078 | 0.92654589625078 | 0.92654589625078 | COX6B1 | 1 | Metabolism |
