## Supplementary material for "Differential Adaptation and Remodeling of the Tumor Microenvironment in Breast Cancer Cells During the Stages of Brain Metastasis": Results of GSVA enrichment analysis performed on the scRNA-seq data, which were grouped based on cell sources (proliferating cells and dormant cells).

### Supplementary Table 4. Results of GSVA enrichment analysis performed on the scRNA-seq data, which were grouped based on cell sources (proliferating cells and dormant cells).

| **Pathway** | Proliferating cells | Dormant cells |
| --- | --- | --- |
| Tumor_Inflammation_Signature | -0.171262175 | 0.188187818 |
| Cellular_response_to_hypoxia | 0.043667379 | -0.238397916 |
| Tumor_proliferation_signature | 0.58832394 | -0.572656497 |
| EMT_markers | -0.401735153 | 0.239331599 |
| ECM-relatted_genes | 0.241610223 | 0.479810903 |
| Angiogenesis | -0.267314132 | 0.3386892 |
| Apoptosis | -0.11297245 | 0.168013017 |
| DNA_repair | 0.071036527 | -0.212751478 |
| G2M_checkpoint | 0.340764695 | -0.319396773 |
| Inflammatory_response | -0.175248928 | 0.261205576 |
| PI3K_AKT_mTOR_pathway | -0.093371947 | 0.028171619 |
| P53_pathway | -0.150952285 | 0.145450729 |
| MYC_targets | 0.399708948 | -0.332056885 |
| TGFB | -0.273439175 | 0.324281897 |
| IL-10_Anti-inflammatory_Signaling_Pathway | -0.169261852 | 0.090334055 |
| Genes_up-regulated_by_reactive_oxigen_species_(ROS) | -0.16167908 | -0.246458347 |
| DNA_replication | 0.490278057 | -0.41557711 |
| Collagen_formation | -0.047701567 | 0.272898204 |
