## Supplementary material for "Differential Adaptation and Remodeling of the Tumor Microenvironment in Breast Cancer Cells During the Stages of Brain Metastasis": Results of GSVA enrichment analysis performed on the scRNA-seq data, which were grouped based on cell cluster.

Supplementary Table 5. Results of GSVA enrichment analysis performed on the scRNA-seq data, which were grouped based on cell cluster.

| **Pathway** | Cluster 0 | Cluster 1 | Cluster 2 | Cluster 3 | Cluster 4 | Cluster 5 | Cluster 6 | Cluster 7 | Cluster 8 |
| --- | --- | --- | --- | --- | --- | --- | --- | --- | --- |
| Tumor_Inflammation_Signature | 0.54676056 | -0.338551251 | -0.008813567 | -0.144436135 | -0.43409071 | -0.465514732 | 0.377117111 | 0.407875666 | 0.27422269 |
| Cellular_response_to_hypoxia | -0.512339305 | -0.01123549 | -0.219382877 | -0.375427307 | -0.033046406 | -0.599587314 | -0.513501406 | 0.526527822 | 0.547741163 |
| Tumor_proliferation_signature | -0.734016994 | 0.519660123 | -0.501843057 | 0.565156788 | 0.817159555 | -0.559588692 | -0.817878307 | 0.600007042 | 0.337584868 |
| EMT_markers | 0.481454652 | -0.653742586 | 0.389911883 | -0.379081051 | -0.006979076 | -0.222245084 | 0.534978208 | 0.16114354 | -0.403797409 |
| ECM-relatted_genes | 0.374381881 | -0.373334688 | -0.117820471 | -0.319178018 | -0.208389733 | -0.018021246 | 0.646471191 | 0.516757072 | -0.030823717 |
| Angiogenesis | 0.335493002 | -0.397246225 | 0.037512992 | 0.007203761 | -0.2120972 | -0.451278246 | 0.342773985 | -0.093072073 | 0.227725151 |
| Apoptosis | 0.129260705 | -0.225603079 | -0.071524893 | -0.148829091 | -0.173389483 | -0.236074377 | 0.070386482 | 0.193278215 | -0.010308832 |
| DNA_repair | -0.377316861 | 0.183105157 | -0.134932932 | 0.26841695 | 0.350281245 | -0.355544586 | -0.42350959 | 0.342410935 | -0.260398269 |
| G2M_checkpoint | -0.628887802 | 0.338814849 | -0.415572489 | 0.458690478 | 0.521634824 | -0.417061712 | -0.561271688 | 0.347817951 | 0.310385503 |
| Inflammatory_response | 0.1713642 | -0.110219075 | 0.055921183 | -0.012711872 | -0.330013383 | -0.190305768 | 0.229234602 | 0.025063233 | 0.041553451 |
| PI3K_AKT_mTOR_pathway | -0.43743092 | 0.07715685 | 0.056810542 | -0.213011894 | 0.18055104 | -0.371944256 | -0.237991786 | 0.183421782 | 0.045860957 |
| P53_pathway | 0.18433204 | -0.136844399 | -0.11890501 | -0.233742992 | -0.044421871 | 0.151867126 | 0.152865893 | -0.006581697 | -0.168633898 |
| MYC_targets | -0.637180968 | 0.471487599 | -0.348102384 | 0.272592288 | 0.275956277 | -0.256917665 | -0.646002641 | 0.542697204 | 0.030525086 |
| TGFB | -0.218805819 | -0.058958825 | 0.09467468 | -0.393584042 | -0.085350132 | -0.406402041 | 0.328823287 | -0.16391675 | 0.278328264 |
| IL-10_Anti-inflammatory_Signaling_Pathway | 0.417137037 | -0.257100028 | -0.255167114 | -0.134664092 | -0.386772879 | -0.463225586 | 0.29368834 | 0.112060086 | 0.022214087 |
| Genes_up-regulated_by_reactive_oxigen_species_(ROS) | -0.310126804 | -0.182159922 | 0.161244084 | -0.05963265 | -0.164564716 | -0.155397178 | -0.325861899 | 0.352551329 | -0.230383964 |
| DNA_replication | -0.628876071 | 0.388242924 | -0.442511135 | 0.773173753 | 0.143524844 | -0.485786532 | -0.589473738 | 0.600888266 | -0.122014971 |
| Collagen_formation | 0.24917257 | -0.201314683 | 0.478068128 | -0.033292505 | -0.216707098 | -0.434971964 | 0.344169793 | -0.11055777 | -0.010305336 |
